# Belowground competition can influence the evolution of root traits

**DOI:** 10.1101/553115

**Authors:** Sara M. Colom, Regina S. Baucom

## Abstract

Although root traits play a critical role in mediating plant-plant interactions and resource acquisition from the soil environment, research examining if and how belowground competition can influence the evolution of root traits remains largely unexplored. Here we examine the potential that root traits may evolve as a target of selection from interspecific competition using *Ipomoea purpurea* and *I. hederacea*, two closely related morning glory species that commonly co-occur in the United States. We show that belowground competitive interactions between the two species can alter the pattern of selection on root traits in each species. Specifically, competition with *I. purpurea* changes the pattern of selection on root angle in *I. hederacea*, and competitive interactions with *I. hederacea* changes the pattern of selection on root size in *I. purpurea*. However, we did not uncover evidence that intraspecific competition altered the pattern of selection on any root traits within *I. hederacea*. Overall, our results suggest that belowground competition between closely related species can influence the phenotypic evolution of root traits in natural populations. Our findings provide a microevolutionary perspective of how competitive belowground interactions may impact plant fitness, potentially leading to patterns of plant community structure.

## Introduction

One of the key reasons plant species are thought to coexist in a given habitat is through a niche partitioning (Aarssen 1997; Huston 1997; Tilman et al. 1997b, 2001; Loreau 2000). Such niche partitioning is hypothesized to occur following competitive exclusion (competitive exclusion under limiting similarity; Gause 1936; Hutchinson 1957; Hardin 1960; MacArthur and Levins 1967), or from trait divergence stemming from competitive interactions between species (*i.e.* character displacement; Brown and Wilson 1965; Pfennig and Pfennig 2009). Because of the relevance of these ideas to the formation of plant community structure, there is a large body of literature examining competitive interactions among plants (Faget et al. 2013). Most of this work, however, focuses on above-ground interactions, and as a result, little is known about root-root interactions belowground.

Roots, which provide a vital resource acquisition function for the plant (Fitter, 2002), also serve as a structure through which plants experience competitive interactions with neighboring plants, whether indirectly through alterations of the soil environment—*i.e.* reduction of water and nutrients—or directly by the excretion of signaling and/or allelopathic molecules (*reviewed in* Shenk, 2006 & Callaway, 2002). The plant root system can be roughly characterized into both size and architectural traits. Root surface area, width, and root length are size proxies whereas traits describing the spatial organization the root system, such as root angle, lateral root branching pattern, and internode branching distance are root architecture traits (Fitter et al. 1991; Lynch, 1995). These root phenotypes strongly influence how a plant accesses nutrients and water (Fitter et al. 1991; BassiriRad, 2005; Manschadi et al. 2006; *reviewed in* Lynch 2007; Kellermeier and Amtmann, 2013). For example, shallow root architectures are linked to increases in the uptake of immobile resources such as phosphorus (Lynch and Brown, 2001; Fitter et al., 2002; Beebe et al. 2006; Lambers et al. 2006), whereas deep root architectures are linked to an increase in water uptake (Beebe et al. 2006; Ho et al. 2005). Thus, shallow root systems may be more advantageous and lead to higher fitness in nutrient-limited soils, and deeper root systems may provide a fitness advantage in water limited environments. How different root traits may influence fitness in the field is most often studied in crop plants (Lynch 2007), leading to a significant gap in our understanding of the factors that influence root trait evolution in natural plant populations. In light of this, how belowground interactions between competing species can mediate plant resource acquisition patterns and potentially alter selection on root architecture and size traits—especially in wild plant species—is largely unknown.

There are plausible reasons to expect competitive interactions to influence the adaptive evolution of root traits. The intensity of competition between plants is greater when rooting zones overlap (*reviewed in* Casper and Jackson, 1997; Casper et al. 2003; Rubio et al. 2003), indicating that occupying the same belowground niche has a deleterious effect on plant fitness. Although studies hint that differences in root systems between competitors leads to higher fitness (*reviewed in* Silvertown, 2014), most of the research characterizes the root system at a coarse level (*e.g.* belowground biomass, root length density; Poorter and Ryser, 2015), and has yet to include specific root size and architectural traits. Competition between co-occurring, closely related species can be especially intense due to greater overlap in physical space or niche use (MacArthur and Levins, 1967; Pfennig and Pfennig 2009; Burns and Strauss, 2011). To reduce the effects of this competition, selection would be expected to favor the divergence of root traits (*e.g*., root angle, root length, and overall root system size) that play important roles in water and nutrient acquisition.

Despite this expectation, there are other explanations for particular root trait shapes or sizes in a species. As above, roots may evolve shallower, deeper, or larger root systems (among other changes) to optimize resource uptake in particular soil environments (Manschadi et al. 2006; Ferguson et al. 2016). Thus, specific root traits may reflect responses to factors in the environment that are distinct from competition. The only way to differentiate competition from other environmental factors that influence root trait evolution is to manipulate the presence of the competitor under otherwise identical conditions and determine if the pattern of selection on the trait is altered as a result (Wade and Kalisz, 1990; Dudley, 1996; Mauricio and Rausher, 1997). While there are many studies assessing how competition influences plant fitness (*reviewed in* Casper and Jackson, 1997 and Faget et al. 2013), there are no studies, to our knowledge, that have examined the potential that competition from a closely related species acts as an agent of selection on root morphology.

The purpose of this work is to determine if belowground competition between two morning glory species—*Ipomoea purpurea* and *I. hederacea*—can influence the phenotypic evolution of root traits in either species. *I. purpurea* and *I. hederacea* are two closely related vines and are common weeds of agriculture in the southeastern and Midwest US. They are most commonly found growing naturally in agricultural fields or in areas of high disturbance (Baucom et al. 2011). In some fields, both species are found to co-occur and intensely compete by vining together above ground; in other fields only one of the species may be present (personal observation, RS Baucom). Previous work has established that competition from one species can alter the pattern of selection on the other. Smith and Rausher (2008) manipulated the presence of *I. purpurea* and experimentally showed that competition between the two species for pollinators can lead to divergence in the floral morphology of *I. hederacea*. Because these species interact in other ways, and share similar morphology as well as resource needs, it is likely that other competitive interactions between the two can lead to trait divergence—namely, root trait divergence following belowground competitive interactions.

Here, we examine the potential that competitive interactions between these two closely related species can drive the evolution of root traits, and we do so by addressing some of the criteria for demonstrating the process of character displacement (*detailed in* Schluter and McPhail, 1992). We first characterize the extent of phenotypic overlap in early growth root traits between *I. purpurea* and *I. hederacea* to determine if the species overlap in the same below-ground niche and then examine the potential for genetic variation underlying these traits. We likewise investigate the potential that natural selection can drive the evolution of root traits in field conditions. We specifically asked the following questions: How do root traits vary within and between species, and to what extent do the species exhibit phenotypic overlap? Is there evidence for genetic variation underlying root traits of either species, indicating that traits can respond to selection? Does belowground interspecific competition between *I. purpurea* and *I. hederacea* impose selection on root traits, and is there evidence that within-species competition (specifically, *I. hederacea-I. hederacea* competition) similarly acts as an agent of selection? Because the adaptive potential of traits can be obscured by plasticity when in competition, we also examine the potential that the presence of a competitor can directly impact root phenotypes. To our knowledge, this is the first study to explicitly test the potential that root traits may exhibit evidence of selection as a result of competitive interactions.

## Materials and methods

### Study system

The common morning glory, *Ipomoea purpurea* (L.) Roth (Convolvulaceae) and ivy leaf morning glory, *I. hederacea* (L.) Jacquin are self-compatible annual climbing vines that commonly co-occur throughout the eastern United States. The two closely related sister species occur in similar habitat types (*e.g*. side of train tracks, agricultural fields, road sides and waste areas). Both species germinate between the months of May and August and begin to flower about six weeks after germination and continue to flower until they die at first frost. The species have similar above-ground growth patterns and produce long stems that branch occasionally. *I. purpurea* is larger (up to 3 m long) compared to *I. hederacea* (up to 1.82 m long). Belowground, *I. purpurea* and *I. hederacea* have fibrous root systems consisting of a primary root with branched lateral roots, and both species vary greatly in the degree of lateral root branching (*personal observation*; *see* fig. 1).

**Figure 1.**
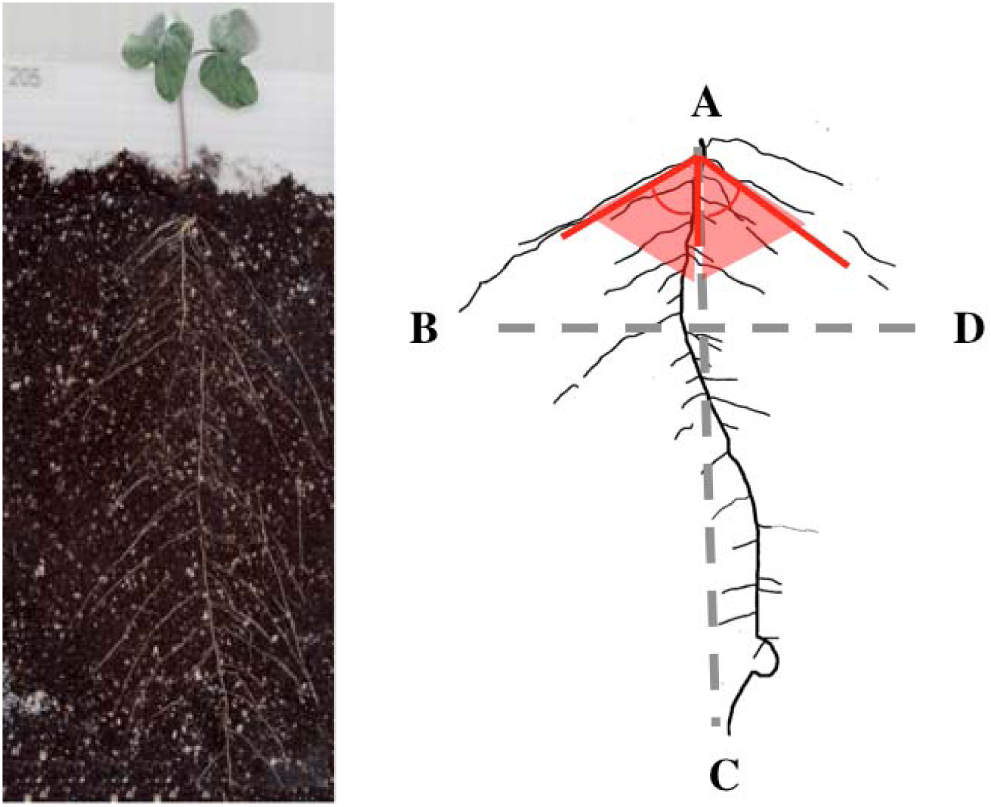
Example of an *Ipomoea* individual growing in rhizotron containing soil (left) and root traced in photoshop (right). Landmarks are placed to estimate root system width (distance between landmark B and D), primary root length (vertical distance between landmark A and C) and left and right root angle (inner angle formed between A, C with respect to B and D, separately for the left and and right root angles, respectively). The grey dashed line indicates the distance calculated for root system width and primary root length, and the red shaded region and solid red arcs indicate the root angle estimated for left and right angle, respectively.

The history of coexistence between *I. purpurea* and *I. hederacea* is only partially known. Evidence suggests that *I. purpurea* is native to Central America (Gray 1886; Barkley 1986; Hickman 1993, Fang, 2013), and it has been present in the eastern United States since at least the early 1700s (Pursh 1814). In contrast, *I. hederacea* has been in the United States for at least 150 years (Bright 1998) but whether *I. hederacea* is native to the United States (Mohr 1901; Stevens 1948) or was introduced from tropical America is uncertain (Shreve et al. 1910; Strausbaugh and Core 1964; Wunderline 1982; Mahler 1984).

### Plant material and growth conditions

We performed complementary greenhouse and field studies to investigate the potential that root traits of these two sister *Ipomoea* species could respond to natural selection. To generate experimental seeds for our common garden and field experiments we selfed 25 and 35 maternal lines of *I. purpurea* and *I. hederacea*, respectively, which were previously sampled as seed from five populations located in Pennsylvania and Ohio. Seeds were scarified and planted in a randomized design in the Matthaei Botanical Gardens (Ann Arbor, MI) greenhouse in November of 2015 and plants were allowed to set seed from selfing for all subsequent experiments.

## Greenhouse experiment

We performed a greenhouse experiment to characterize early growth root traits between and within *I. purpurea* and *I. hederacea* in the summer of 2016. We planted replicate once-selfed seeds in custom built rhizotrons containing generic potting soil (fig. 1) in greenhouse conditions at the Matthaei Botanical Gardens (Ann Arbor, MI). Rhizotrons consisted of 20.32 cm × 25.4 cm frames made out of cut pieces of corrugated plastic and a transparent polystyrene sheet held to the frame by duct tape. Each rhizotron was filled with 20.45 grams of soil and a single seed was placed in the center of the rhizotron approximately one inch below the soil surface against the transparent polystyrene sheet.

We planted three replicates per maternal line per species in the rhizotrons and positioned the rhizotrons in custom-built wooden frames at 30° in a completely randomized design (see fig. A1 in appendix A for root image from rhizotron, and instructions on building rhizotron frames in app. fig. B2; both apps. A and B are available online). We replicated this experiment in its entirety, twice. Thus, we planted 150 individuals of *I. purpurea* (3 biological replicates *x* 25 maternal lines *x* 2 experimental replicates) and 210 individuals of *I. hederacea* (3 biological replicates *x* 35 maternal lines *x* 2 experimental replicates) for a total of 360 individuals. We watered each individual daily by hand to standardize water availability across all individuals for three weeks.

### Greenhouse root phenotyping

Two weeks after germination, we scanned each rhizotron to measure below-ground root traits using a CanoScan LIDE 110^®^ scanner bed. For each image, we traced the roots in Photoshop version CS6 to facilitate automated quantification of root size based on their pixels in ImageJ version 3.0 (Abràmoff and Magalhães 2004).

We focused on root size and root architecture by measuring root system pixels (root size), root system width, primary root length and root angle on the two week old seedlings (fig. 1). We elected to focus on these specific traits because they are relatively straightforward to measure across different growing conditions and they also play a vital role in plant resource use and uptake (Wasson et al. 2012; Paez-Garcia, et al. 2015). Prior to data collection, ImageJ was first globally calibrated with the set scale tool in order to obtain measurements in metric units for all of the following procedures. To obtain primary root length, root system width and root angle, first we used the multi-pointer tool and placed a total of four points along the root tips of the root system image in the following order: 1) primary root at the root stem surface, 2) root tip of the left outermost root, 3) primary root tip and 4) root tip of the right outermost root.

We used the statistical programming language R (R Core Team, 2017) to calculate primary root length, root system width and root angle (the script is available in GitHub at https://github.com/SaraMColom/Selection_RootTraits_2016_2017). For primary root length, we calculated the vertical distance between the primary root at the soil surface and the tip of the primary root. We estimated root system width as the euclidean distance between the outermost lateral root tips. To estimate root angle (θ) we used the cosine formula,

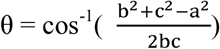

on a right triangle formed by the primary root (the longest root perpendicular to the soil surface), and each of the outer lateral roots. Here, *b* is the distance between the primary root and the outermost lateral root tip, *c* is a measure of the length of the primary root calculated above, and *a* is the length between the outermost lateral root tip and the primary root tip (fig. 1). Root angle was calculated for both the right and left lateral roots, separately averaged, and reported in degrees. We elected to use the angle made between the primary root and the outermost left and right lateral roots because previous research has shown that this trait is indicative of root architecture types (Lynch and Brown, 2001; Uga et al. 2013; Colombi et al. 2015). Finally, to obtain root size, we converted the traced root images into binary images, selected ‘area’ as a measurement output in the ‘measurement’ option, and performed the ‘Analyze Particles’ function in ImageJ. This function quantified the total number of black pixels (all pixels from the root system) and reported the values in centimeters squared.

## Field experiment

### Field design and planting

We conducted a field experiment in the summer of 2017 to characterize root trait variation in *I. purpurea* and *I. hederacea* grown in field conditions and to determine if natural selection acts on root traits in the context of interspecific competition. We planted replicate selfed seed from maternal lines of *I. purpurea* and *I. hederacea* in two treatments: ‘alone’and interspecific competition (fig. 2) in a field plot at Matthaei Botanical Gardens (Ann Arbor, MI) on June 2, 2017. We likewise planted replicate selfed seed from *I. hederacea* maternal lines in the presence of intraspecific competition to determine if, at least for this species, within-species competition could influence the evolution of root traits. We used eight maternal lines of *I. purpurea* and *I. hederacea* from a single population from Pennsylvania (PA4). We decided to use maternal lines from this population since preliminary greenhouse data demonstrated high phenotypic overlap between both species for this population (fig. B1). We did not fertilize the field beforehand, nor is it land that has crop rotation. We planted eight replicates of each maternal line per species across four blocks for our alone treatment (8 maternal lines *x* 2 biological replicates *x* 4 blocks *x* 2 species = 128 plants). For our interspecific competition treatment we paired each maternal line of *I. purpurea* with each maternal line of *I. hederacea* for each possible pairwise combination, and planted 2 replicates of each pairing across 4 blocks (64 unique interspecific competition combinations *x* 2 biological replicates *x* 4 blocks = 512 individuals *x* 2 species = 1024 plants). For our intraspecific competition treatment we planted 2 replicates of each unique combination of the 8 *I. hederacea* maternal lines within each block (28 unique interspecific competition combinations *x* 2 biological replicates *x* 4 blocks = 224 experimental units *x* 2 plants = 448 plants). Each pairwise competition pairing was replicated 8 times across the experiment. It is important to note that although we were likewise interested in the potential that *intraspecific* competition in *I. purpurea* could influence root trait evolution in this species, we elected to examine this only in *I. hederacea* due to both field space limitations and the experimental difficulty of phenotyping large numbers of root systems in the field.

**Figure 2.**
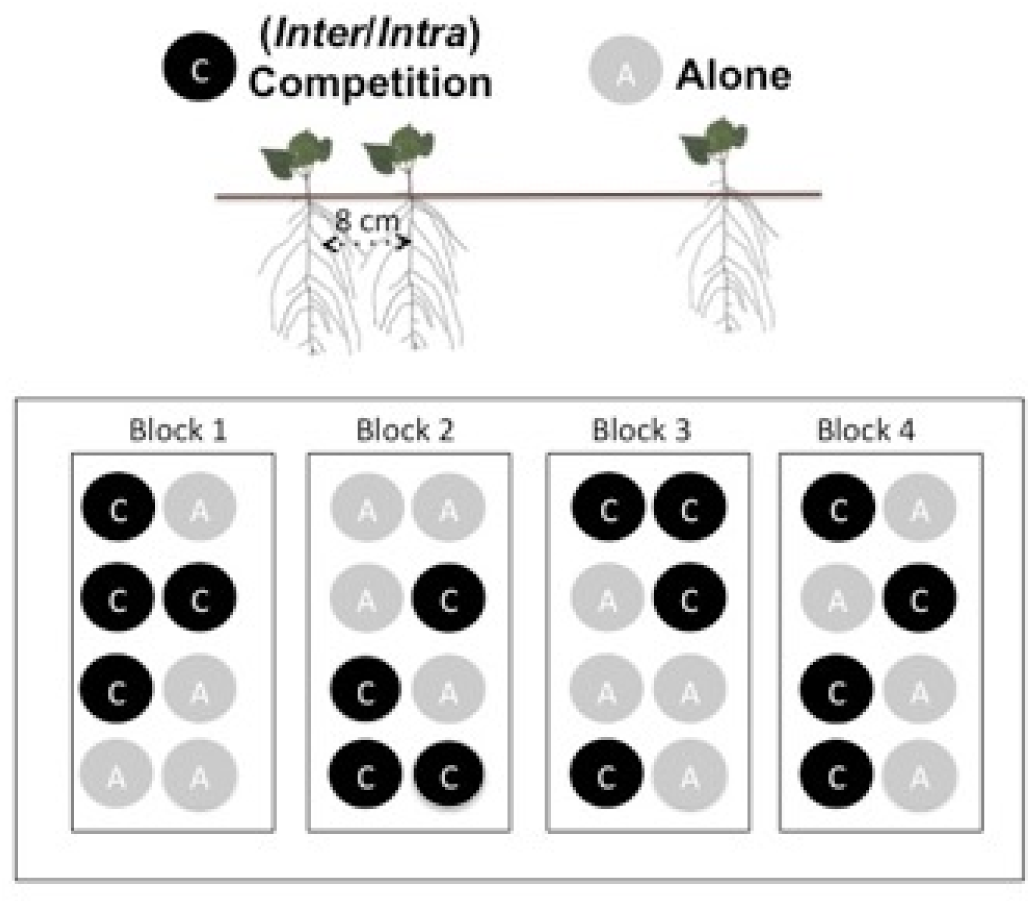
Diagram of the field experiment showing *Ipomoea* plants grown in the presence of competition (inter- or intraspecific) or alone. Competition treatments (inter- or intraspecific) are indicated by solid black circles with a ‘C’, and the alone treatment is indicated by a gray solid circle with an ‘A’. Each experimental unit (*i.e.* each unique competition pairing and alone treatments) was replicated eight times and randomly arrayed with two biological replicates per experimental unit per block.

Seeds were planted into experimental units (*i.e.* cell with either a single plant for the absence of competition, or two plants for the competition environment) which were arrayed across the four spatial blocks in a completely random block design. Experimental units were spaced uniformly 1m from each other; individuals in competition treatments were planted 8 cm apart from each other within their experimental unit. One week after planting we scored germination. Due to intense drought < 48% of seeds germinated overall. The average precipitation for June in 2017 in Ann Arbor, MI was 0.10 cm. In comparison, the average preciptation was 0.18 cm and 0.20 cm in 2016 and 2018, respectively based on data collected online from National centers for Environmental Information (NCEI; Menne et al. 2012). We thus planted a second experimental cohort (cohort 2) on June 19, 2017 to increase sample size, and this cohort was planted to conserve the same level of replication across all experimental units. We had 86 % germination success with replanted individuals, and ended with a total 1177 plants of which 670 plants were in interspecific competition, 341 plants were in intraspecific competition, and 166 plants in the alone treatment; of our final sample, 56% came from cohort 1 and 44% came from cohort 2. Throughout the timespan of the field experiment we kept the immediate surroundings of each experimental unit (∼15 cm from base of plants) clear of weeds. Three weeks after the first planting date we placed 1m tall bamboo stakes at the base of every experimental plant at a 45° angle, which allowed us to train vines of competing plants away from another, thus removing the potential for above-ground competition.

### Field root excavation

To characterize the phenotypic variation of root traits of *I. purpurea* and *I. hederacea* grown in field conditions with and without competition, we adapted the ‘shovelomics’ excavation method described by Colombi et al. (2015). We harvested roots after three weeks of growth on a subset of plants in the field. For root phenotyping, we sampled individuals only from cohort 1 because individuals from cohort 2 were small and not reproductively mature, whereas most individuals of cohort 1 had developed flower buds. We sampled between two to four replicates per maternal line from both competition treatments—specifically, we sampled a total of 165 *I. purpurea* individuals, (N = 23 and N = 142 from the alone and interspecific competition treatments, respectively), and a total of 304 *I. hederacea* individuals (N = 31, N = 132, N = 141 from alone, interspecific and intraspecific competition treatments, respectively). To excavate roots, we cut the stem 5 cm from the soil surface, marked the side of the stem facing the competitor with a permanent marker and then dug the root system with a shovel by placing the shovel head at 45 degree angle, 15 cm from the plant stem. This method unearthed the first 15 cm of the root system. The excavated root core was then shaken gently to remove adhering soil and placed in plastic bags for root imaging.

### Field root imaging and phenotyping

Root imaging was carried out indoors with the use of a cubic photo shooting tent (MVPOWER, 40 cm × 40 cm × 40 cm) in order to standardize imaging between samples and facilitate the use of REST (Colombi et al. 2015), an automated root phenotyping program developed to characterize the root systems of plants grown in the field. To image the root system, roots were hung in the center of the photo shooting box and photographed with a Canon EOS Rebel XSi 12.2 MP (18-55 mm IS Lens). Images were imported into REST (Colombi et al. 2015), and we manually specified where the stem at the soil surface was for each image (shown in fig. 3). After user specification of the stem/soil surface, REST draws a rectangular region of interest for pixel analysis to standardize measurements across images. All root measurements were quantified from the pixels lying within this region of interest. REST returned the root angle (right and left root angle), root system width (‘Max width’) and a root system size proxy (‘Area convex hull’), among other morphological and architectural traits. We focused on these three root traits because they are similar to the traits captured in our greenhouse study. Root angle (left and right) are determined by calculating the outermost angle between the top most lateral root and the soil surface plane at the plant stem, and then subtracting this value from a perpendicular line (*i.e.* 90°) drawn at the plant stem. The root system width captured in REST is the same measurement as taken in the greenhouse rhizotron study as they were both estimated as the euclidean distance between the root tips of the left and right outermost lateral roots. In contrast, root system size estimated from the greenhouse rhizotron study and REST program were similar, but not identical. Root size from the greenhouse rhizotron study was based on the total area of root derived pixels, and root size in REST was based on the convex hull of all root derived pixels. We do not have data for primary root length from plants grown in the field because this trait was destroyed in the process of sampling the roots.

**Figure 3.**
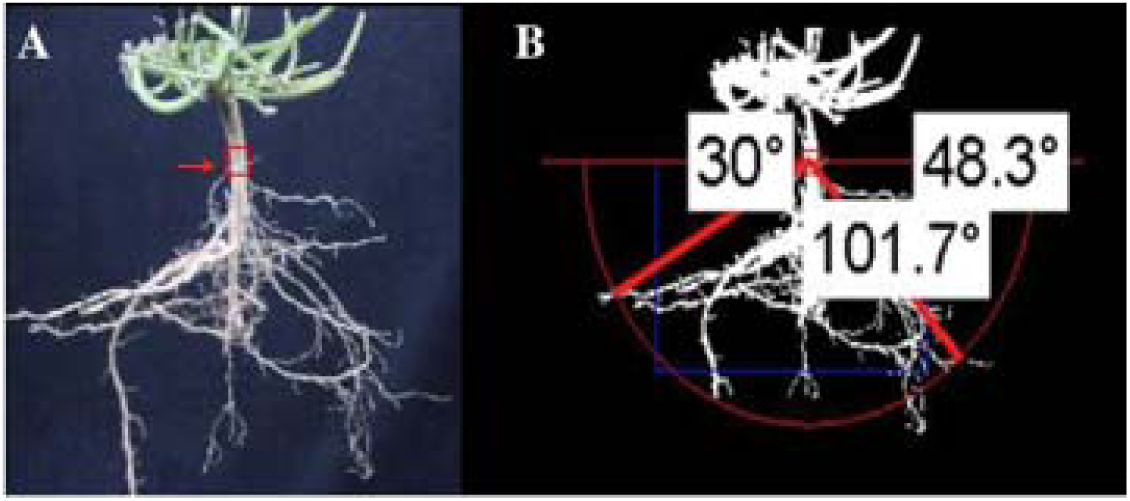
REST output showing the original image of an *Ipomoea* species excavated from the field (A) and its binary form with an arc superimposed by REST to obtain the outer right and left root angles (here 30 degrees on the left and 48.3 degrees on the right) from the horizontal place in red. (B) The blue box shows the region of interest detected automatically by REST program of which total root based pixels were quantified from within to obtain root size (‘area convex hull’ in REST) and measure root system width (‘max root width’ in REST) based on the distance between the right and left outermost roots in the box.

### Field plant fitness data

We began to collect mature fruit in the month of September and continued to do so until late October when all plants have senesced. The entire aboveground of all remaining plants were collected by first frost and seeds were manually removed, cleaned, and counted with a seed counter to obtain an estimate of total fitness. We sampled 165 individuals of *I. purpurea* (N = 23 and N = 142, from alone and competition treatments, respectively), and 304 individuals of *I. hederacea* (N = 31, N = 132, and N = 141, from alone, interspecific and intraspecific competition treatments, respectively). Ultimately, we sampled seeds from a total of 508 plants; 27% of these individuals came from cohort 1 and 72% from cohort 2.

## Data analysis

### Greenhouse experiment

All statistical analyses were carried out in R (version 3.3.1). We fit separate linear mixed models using the ‘lmer’ function of the lme4 package (Bates et al. 2015) for each of the root traits measured to test for the presence of variation in root traits between species, populations, and maternal lines. Each respective linear mixed model consisted of the root trait as the response variable, species, population and experimental replicate (*i.e.* temporal replicate; ‘Experiment’) as fixed effects and maternal line as a random effect; *i.e.* Root trait ∼ Experiment + Population + Species + (1|Population: Maternal line) + *ε*. To ascertain the significance of the predictor variables we used F-statistics for the fixed effects, with Satterthwaite’s method to estimate denominator degrees of freedom, and a log likelihood ratio test to estimate chi statistic (χ^2^) for the random effect (using the ‘anova’ and ‘ranova’ functions from the lmerTest package; Kuznetsova et al. 2017). We ran additional linear mixed models for each species separately to test for evidence of maternal line variation within *I. purpurea* and *I. hederacea*, where root trait was the response variable, population and experimental replicate were fixed effects and maternal line was a random effect. We further examined how roots varied between species in trait space by performing principal component analysis (PCA) using a correlation matrix of all root traits measured in the greenhouse with the ‘PCA’ function from the FactoMineRPackage (Lê et al, 2008).

### Field experiment

To examine how root traits vary between *I. purpurea* and *I. hederacea* grown in field conditions, and to determine if root phenotypes were influenced by competitive interactions, we ran mixed linear models as above. We fit a separate model for each root trait where the trait was the response variable and block, treatment, and block *x* treatment interaction were fixed effects and maternal line and maternal line *x* treatment interaction were random effects, *i.e.* Root trait ∼ Block + Treatment Block × Treatment + (1|Maternal line) + *ε*. Preliminary analyses indicated that there were no significant maternal line *x* treatment interactions for any trait. We thus elected to exclude these effects from our final models. As above, we visualized phenotypic variation in root traits between species when grown in field conditions by performing principal component analysis (PCA) with a correlation matrix on all traits including root system size, root system width and average root angle. In addition, we generated a correlation matrix using the family mean values for all the root traits measured for each species separately to examine relationships between the three traits.

### Selection analyses

We used genotypic selection analyses (Lande and Arnold, 1983; Rausher 1992) to estimate selection gradients on each root trait in each competition environment, and ANCOVA to determine if competition and experimental block altered selection on root traits of the two species. We elected to perform a joint selection analysis using maternal line averages of the root traits because it allowed us to examine direct selection acting on each trait while controlling for environmentally induced biases (Rausher 1992). The maternal lines were averaged by species, treatment and cohort for our selection gradient analysis. We estimated selection gradients on root system width, root system size, and root angle of both species in each competitive treatment environment (alone and, interspecific and intraspecific competition) by performing multiple regression with the focal root traits included as predictor variables and relative fitness (total seed number divided by the mean seed number by species and treatment) as the dependent variable. For all selection analysis, we used mean standardized root trait values and untransformed relative fitness. Preliminary analysis indicated that individuals of both species from the second cohort produced significantly fewer total seeds than individuals from the first cohort (*I. purpurea*: F_1_ = 100.3, p-value < 0.001; *I. hederacea*: F_1_ = 213.9, p-value < 0.001), but preliminary analyses also provided no evidence that selection gradients differed between cohorts within either species for any root trait. Thus we elected to combine cohorts in the genotypic selection analyses (cohort 1 N = 141 and cohort 2 N = 367). Further, while we examined the potential for non-linear selection influencing root traits in preliminary analyses, we did not find evidence of either stabilizing or disruptive selection acting and thus present only the results of linear selection analyses.

We used analysis of covariance (ANCOVA) to determine if the direction and/or intensity of selection varied between the presence and absence of competition (Wade and Kalisz, 1990) separately for each species. For *I. purpurea*, we compared selection gradients between plants grown in interspecific competition or grown alone, and for *I. hederacea*, we compared selection gradients from inter- or intraspecific competition with that of plants grown alone. In each analysis, models included competition treatment, block, the standardized root trait values, and all interactions as predictors of relative fitness. Significant interactions between the competition treatment and standardized root traits indicate that selection gradients differed between treatments. Block and block *x* treatment interactions were likewise included within the ANCOVAs.

## Results

### Greenhouse experiment

In our greenhouse rhizotron study assessing early root traits, we found significant variation between species in root system width and average root angle (table 1). The root system of *I. hederacea* was wider (8.49 cm, table 1) than that of *I. purpurea* (6.92 cm, table 1) and the overall root size of *I. hederacea* was on average greater (3.95 cm^2^, table 1) than *I. purpurea* (2.37 cm^2^, table 1). *I. purpurea* exhibited lateral roots that were closer to the soil surface (root angle: 37.33 degrees on average, table 1) compared to *I. hederacea* (30.44 degrees on average, table 1). Although species varied in the above traits (table 1), a visualization of the four root traits in a PCA identified considerable overlap of root phenotypes between species (fig. B1*A*). The root system width, root angle and primary root length loaded most strongly on the first principal component, which captured 37.3% of the total variation (fig. B1*B*), and root size loaded most strongly on the second principal component, which explained 29.1% of the total variation (fig. B1*C*). These first two PCA’s can thus serve as descriptors of root system architecture (*i.e.* spatial arrangement of root system) and root size, respectively.

**Table 1.** Species differences in *I. purpurea* and *I. hederacea* root traits measured from rhizotrons in the greenhouse. Least-squares means ±1 SE for each trait by species and F-statistics and likelihood ratio test statistics (χ^2^) values showing the effects of experimental replicate, species, population, and maternal line variation on plant phenotypes. Maternal lines were nested within populations.

| Trait | Species | | F-statistics | | | $\chi^2$ |
| --- | --- | --- | --- | --- | --- | --- |
|  | *I. purpurea* | *I. hederacea* | Experiment<br>*df* = 1 | Species<br>*df* = 1 | Population<br>*df* = 4 | Maternal<br>Line<br>*df* = 1 |
| Root system width (cm) | 6.90 ± 0.28 | 8.49 ± 0.22 | **315.79** | **17.18** | **6.82** | **5.10** |
| Primary root length (cm) | 11.30 ± 0.61 | 11.40 ± 0.46 | **22.52** | < 0.01 | 0.76 | **5.22** |
| Root angle (degrees) | 37.40 ± 1.33 | 30.40 ± 1.01 | **61.42** | **15.04** | 0.22 | **12.47** |
| Root system size (cm<sup>2</sup>) | 2.36 ± 0.21 | 3.95 ± 0.16 | **35.22** | **31.41** | **7.29** | 2.22 |

Additionally, we found evidence for both population and maternal line variation in root traits. We found significant population variation for root system width and root system size, and variation among maternal lines for root system width, root angle, and primary root length (table 1). Separate mixed models, performed per species, identified significant maternal line variation within *I. purpurea* for root system width (χ^2^ = 7.46, p-value = 0.01) and root angle (χ^2^ = 4.05, p-value = 0.04), and marginally significant maternal line variation for root size (χ^2^ = 3.63, p-value = 0.06). We identified maternal line variation within *I. hederacea* for root angle (χ^2^ = 8.63, p-value < 0.01), and marginally significant maternal line variation for primary root length (χ^2^ = 3.10, p-value = 0.08).

### Field experiment

A visualization of root system width, size, and root angle in a principal component analysis showed a high overlap between species in root phenotypes (fig. B1) in plants grown in the field. We identified maternal line variation in root system width (table B1); a within species examination revealed this result to be driven by *I. purpurea* (χ^2^ = 4.69, p-value = 0.03). We further found a significant and strong correlation between root size and root width (r = 0.85, p-value < 0.001; table B2) in *I. purpurea*, whereas there was evidence for strong and significant positive correlations between all root traits within *I. hederacea* (root width and root angle r = 0.59; root size and root angle r = 0.60; root size and root width r = 0.80; p-value < 0.001 for all pairwise traits; table B2).

With the exception of a marginally significant treatment effect on root size (F_2_ = 2.33, p-value = 0.10; table B1), we found that interspecific competition in the field did not strongly influence root phenotypes of either species. A closer examination of the linear mixed models within species suggested that this treatment effect likely impacts *I. hederacea* (F_2_ = 2.10, p-value = 0.12) but not *I. purpurea* (F_2_ = 0.04, p-value = 0.84). However, there was a strong effect of competition on fitness, with *I. purpurea* experiencing a fitness reduction of 30.31% and *I. hederacea* a reduction of 36.47% when in interspecific competition. *I. hederacea* planted in intraspecific competition likewise experienced a significant fitness reduction (39.67 % lower than plants grown alone). Intraspecific competition between *I. hederacea* plants led to slightly lower fitness than when grown in interspecific competition (*i.e.* 6.16% reduction in intra-versus interspecific competition), but this difference was not significant (table 2).

**Table 2.** Influence of competitive treatment on *I. purpurea* and *I. hederacea* root traits when grown in the field. Least-squares means ±1 SE for each trait in each treatment.

|  | *I. purpurea* |  | *I. hederacea* |  |  |
| --- | --- | --- | --- | --- | --- |
| Trait | Alone | Competition | Alone | Competition (inter) | Competition (intra) |
| Root system width (cm) | 7.12 ± 0.37 | 7.00 ± 0.25 | 7.43 ± 0.36 | 7.31 ± 0.25 | 7.03 ± 0.25 |
| Root system size (cm<sup>2</sup>) | 77.50 ± 3.94 | 76.69 ± 2.16 | 78.08 ± 3.77 | 77.28 ± 2.22 | 71.07 ± 2.23 |
| Root angle (degrees) | 26.62 ± 2.05 | 24.43 ± 1.14 | 27.00 ± 1.96 | 24.81 ± 1.16 | 22.63 ± 1.17 |
| Seed number | 750.01 ± 75.69 | 491.15 ± 53.88 | 709.84 ± 68.42 | 450.98 ± 53.48 | 428.26 ± 55.10 |

### Selection on root traits in field conditions

From our selection gradient analyses, we identified positive linear selection on root angle in *I. hederacea* in the presence of interspecific competition (β = 0.23, p-value = 0.03; fig. 4; table B3) but no evidence of selection when when grown alone (β = 0.01, p-value = 0.86; table B3). ANCOVA revealed a significant treatment interaction for root angle in *I. hederacea* (F_2_ = 4.37, p-value = 0.04; table B4), supporting the idea that the pattern of selection for root angle differs according to competitive context for this species. Further, we found a marginally significant block *x* treatment interaction (F_11_ = 2.28, p-value = 0.09; fig. 4; table B2) for root angle in *I. hederacea* when in interspecific competition, suggesting that environmental differences can influence the strength and/or direction of selection on this trait. In comparison, we found no evidence of selection when *I. hederacea* was grown in intraspecific competition (β = −0.27, p-value = 0.29; fig. 4; table B3 and table B4).

**Figure 4.**
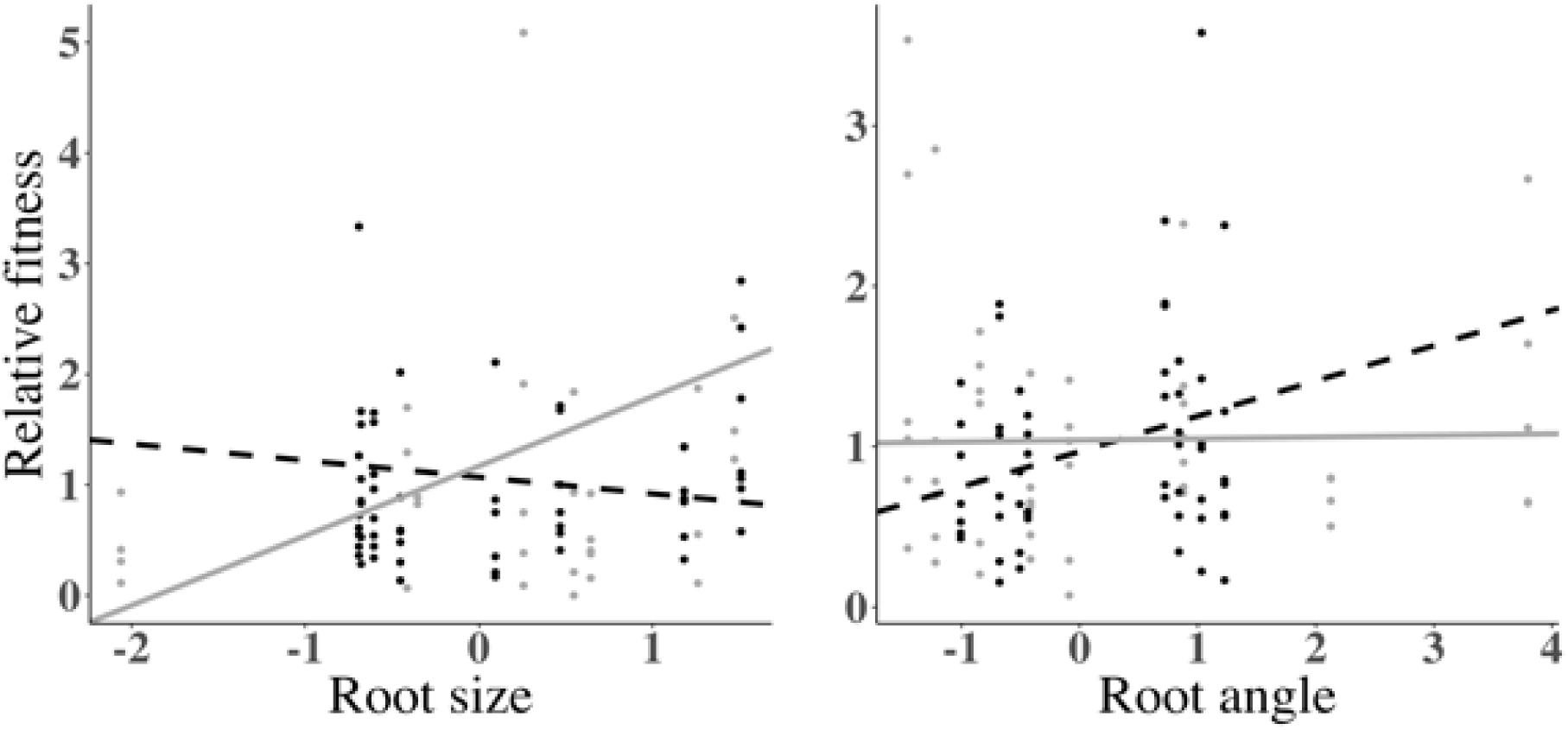
Interspecific belowground competition alters the pattern of selection for root size in *I. purpurea* (A), and root angle in *I. hederacea* (B). Solid grey and solid black circles represent the family mean values of standardized root traits (root size in *I. purpurea* (A) and root angle in *I. hederacea* (B)), with family mean values of relative fitness regressed onto each trait when plants were grown in interspecific competition or grown alone, respectively. Solid grey lines and dashed black lines represent the slope (β) for plants grown in interspecific competition, and plants grown alone.) The β for root size in *I. purpurea* grown alone (0.56 ± 0.29) differed significantly (F_1_=4.88, p-value=0.03; Table B4) from the β of *I. purpurea* grown in interspecific competition (−0.15 ± 0.22). The β for root angle in *I. hederacea* grown alone (0.01 ± 0.08), differed significantly (F_1_=4.37, p-value=0.04; Table B4) from the β of *I. hederacea* grown in interspecific competiton (0.23 ± 0.09).

For *I. purpurea*, we found marginally significant positive selection for root system size when grown in the absence of interspecific competition (β = 0.56, p-value = 0.08; table B3), but no evidence of selection on root size when in the presence of competition (β = −0.15, p-value = 0.51; B). ANCOVA revealed a significant treatment interaction with root size (F_2_ = 4.88, p-value = 0.03; table B4) indicating that selection on root size differs according to competitive context in this species. We likewise found a significant treatment *x* block interaction for root system width within *I. purpurea* (F_11_ = 6.29, p-value <0.01; table B4) and a marginally significant treatment *x* block effect for root angle (F_11_ = 2.20, p-value = 0.10; table B4), indicating that the pattern of selection on these two traits are impacted by both competitive context and other unmeasured environmental variables (*i.e.* block effect).

## Discussion

Given the functional importance of root systems, we hypothesized that competition between two closely related species could impose selection on root traits, and that selection could promote divergence in such traits. However, there are few, if any, examinations of the potential for selection on root traits in field conditions. Thus, we characterized the phenotypic variation in root traits of two closely related species, determined if genetic variation in these traits was present both within the greenhouse and in the field, and examined the potential that competition changed the pattern of selection on roots. From our greenhouse experiment, we found early growth root traits to differ significantly between the species, and found evidence for genetic variation underlying traits—both between population variation and maternal line variation. Results from our manipulative field experiment showed that in the absence of competition there was a trend for positive linear selection acting on root size in *I. purpurea*, but no evidence for the same pattern of selection in the presence of competition. Interestingly, we found selection acting on different traits in *I. hederacea*: in this species, we found positive selection acting on root angle when in the presence of competition with *I. purpurea*, but no evidence of selection on this trait in the absence of competition. Somewhat surprisingly, we found no evidence of selection on root angle (or any root trait) in *I. hederacea* when in the presence of *intraspecific* competition (i.e., *I. hederacea-I. hederacea* competition). Thus, competition below-ground from *I. purpurea* promotes the evolution of broader root angles (*i.e.* a more shallow root system) in *I. hederacea*, but the same effect is not seen in *I. hederacea* when in within-species competition.

Because water, ion and minerals are heterogeneously spread in the soil according to their chemical and physical properties, differences in root architecture between plants determines what specific resources are readily available for uptake and in turn how plants compete for such resources (Lynch, 2005). This provides a likely explanation for the pattern of selection we identified for *I. hederacea* in the presence of interspecific competition; because shallow lateral roots enable the exploitation of nutrients near the soil surface, individuals with shallow roots may be at a fitness advantage when in the presence of a competitor compared to individuals with deeper root systems. In support of this idea, shallow rooting systems have been shown to be advantageous in common bean, maize and rice when grown in environments that are limited by phosphorus and other resources that accumulate in the topsoil (Rubio et al. 2003; Lynch and Brown, 2001; York et al. 2015; Sandhu et al. 2016).

From an ecological standpoint, it is somewhat puzzling that selection favors a larger root system in *I. purpurea* when competition is absent, but not when competition is present. Larger root systems allow for greater exploitation of soil nutrients and water, and have been shown to be correlated with increased fitness in other species (Svačina and Chloupek, 2014; Ehdaie and Waines, 2008). As such, we expected to identify selection for larger root systems regardless of competitive environment. A potential explanation for our findings is that root traits that were not measured here—primary root length, lateral root placement, and/or hair root density—may play an important role in resource uptake in the presence of competition. An investment in greater root foraging precision, as well as selection on traits that optimize resource uptake efficiency could potentially reduce the deleterious effects of competition. Thus, it is possible that root size is not under selection when these two species compete because selection is instead acting on traits that increase resource uptake efficiency (Fitter et al. 1991; Hodge et al. 1999; York et al. 2015).

Although we identified selection on only two traits—root size and angle—the strong correlations we uncovered between root width, size, and root angle suggests traits not under direct selection will likely evolve due to indirect selection. We identified strong positive correlations between root width, size, and angle in *I. hederacea*, indicating that width and size may evolve indirectly given selection on root angle. In *I. purpurea*, the strong positive correlation between root width and size, and pattern of positive selection on root size, suggests that root width should likewise experience indirect positive selection. That we found no evidence of correlations between root angle and root width and size in *I. purpurea* suggests root angle may evolve with fewer constraints in this species. It is likewise notable that we uncovered genetic variation underlying only root width in *I. purpurea* in the field experiment; however, this result is not particularly surprising given that genetic variation in field conditions is often obscured by high environmental variation (Conner, 2003). Notably, in our greenhouse experiment, we found evidence for both population and maternal line variation on root traits in both species, suggesting these traits have the capacity to evolve either through selective pressures or as a result of genetic drift.

Further, while we identified different patterns of selection across the competitive environments between the two species, we found suggestive, but limited, evidence for plasticity in the root traits of either species as a result of competition. Plant root systems can impact the root growth of other closely neighboring plants either indirectly *via* altering the physical and chemical soil environment and/or directly through the excretion of signaling and/or allelopathic molecules (Schenk 2006; reviewed in Cahill and McNickle 2011 & Depuydt, 2014). As such, we expected to find a significant treatment effect on root trait phenotypes, and thus evidence of phenotypic plasticity in root architecture and size traits. Other experiments characterizing root phenotypes in a range of species have found mixed results when plants are grown in competition, ranging from genotypic- and species-specific responses in root growth to no response whatsoever (Mahall and Callaway 1991; Falik et al. 2003; Bartelheimer et al. 2006; Fang et al. 2013; Belter and Cahill 2015; Litav and Harper 1967; Semchenko et al. 2007). Therefore, the results we report here suggest that these two *Ipomoea* species may lack a mechanism to modify their root growth in competition, that plasticity may be occurring in other, unmeasured traits, or simply that the effect sizes on root trait changes due to competition were small, and high variance due to other environmental factors (e.g. potentially the influence of drought in the 2017 field season) reduced our ability to identify significant plasticity in root traits given competition.

Importantly, the belowground plant-plant competition imposed by our experimental design led to reduced fitness of both species—around 35% fewer seeds produced by each species in the presence of competition (whether interspecific and intraspecific)—indicating that although we did not uncover root trait plasticity, there was clearly a cost imposed by the presence of belowground competition between and within species. We note, however, that the strongest trend in reduced root size occurred when *I. hederacea* was planted in *intraspecific* competition relative to interspecific competition. This suggests *I. hederacea* may potentially be decreasing overall plant growth as an adaptive response to reduce intraspecific competition. Such a potential plastic response within *I. hederacea* when in competition with a congener may explain why we did not detect evidence for selection from intraspecific competition on root traits in this species.

Overall, our finding of different patterns of selection acting on root traits in the different competitive treatments indicates that plant-plant competition can act as a selective agent on root traits. That we identified selection on different root traits between species is consistent with the idea of niche partitioning, which predicts greater divergence in resource associated traits between species to reduce competition for limiting resources (MacArthur and Levins 1967). Multiple field studies examining the relationship between different rooting depths of various co-occurring plant species have shown that a decrease in overlap between rooting zones of neighboring plants positively impacts plant yield and biomass (*i.e.* plant fitness; Fargione and Tilman 2005; Mueller et al. 2013). Our results extend this finding to show that interactions between two closely related, co-occurring species elicits selection for different patterns of root traits. Hence, it is possible that competition between the two *Ipomoea* sister species promotes the divergence in resource-related root traits.

Finally, although our research provides the first experimental evidence that belowground competition can influence the evolution of root traits in these two related species, we are not showing the outcome of such competitive interactions across many natural populations. More specifically, while our study supports the idea that the adaptive process can occur in root traits as a response to belowground competition, we do not explicitly test for broad-scale patterns that would suggest such interactions have led to trait divergence (*i.e*., divergence in root traits where the species co-occur versus similarity in areas where they do not co-occur). Future work testing for patterns of phenotypic evolution in root traits between multiple naturally occurring populations of these two species is thus needed to draw conclusions for, if and how competition belowground has influenced the evolution of root traits in natural populations across the landscape.

## Acknowledgements

We thank Corlett Wood for helpful comments on earlier versions of this article. We also thank Tyler Marrs for help with the construction of rhizotrons and wooden frames, and Andres Ibarra, Megan Van Etten, Diego Alvarado-Serreno, Sonal Gupta, Teresa Dorado and Lourdes Abreu Torres for their invaluable assistance with planting, maintenance of the field site and sample collection. We thank Gloria Brana and Gloria Colom for help designing rhizotrons and wooden frames, and employees of Matthaei Botanical Gardens, especially Michael Palmer and Paul Girard for their expertise and loan of field equipment and machinery. This work was made possible with financial support of internal grants at the University of Michigan and Matthaei Botanical Gardens Winifred B. Chase Fellowship.

## Data availability statement

The R code is available at GitHub at https://github.com/SaraMColom/Selection_RootTraits_2016_2017, and the data will be uploaded to the Dryad Digital Repository.

## Appendix A

### Custom made rhizotron frame instructions

**Figure A1.**
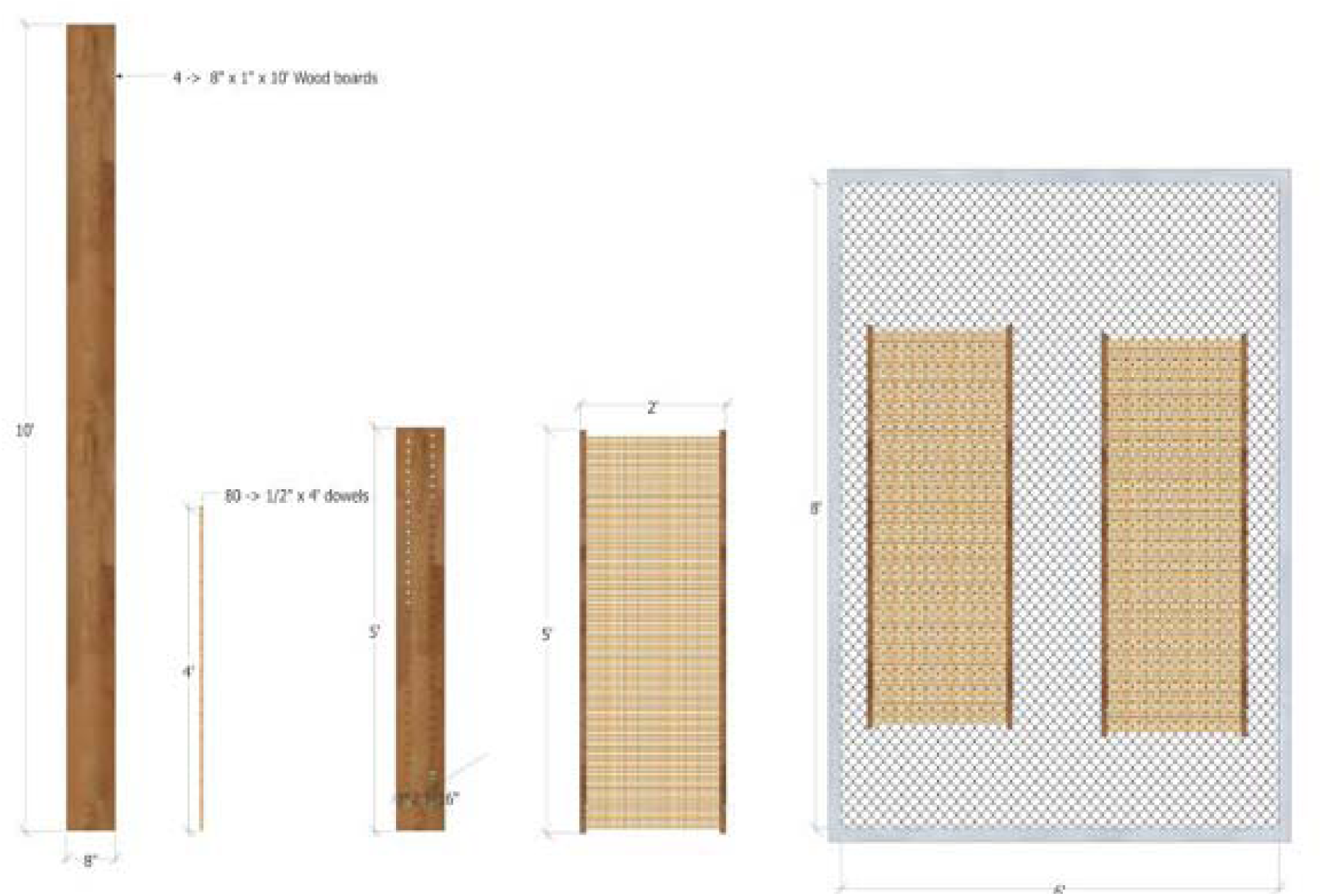
Procedure and materials required for construction of rhizotron frames. Four wood boards of 8” × 1” x 10’, 80 1/2” × 4’ wooden dowels, wood glue, power drill, measuring tape, and a pencil will be required to build a total of four rhizotron frames. First saw each 10’ wooden board in half to obtain a total of eight 5’ boards and then saw each 4’ wooden dowel in half to obtain a total of 160 2’ dowels. Measure and draw out 40 evenly spaced dots on the 10’ board approximately two inches from the top edge. For every of the 40 evenly spaced dots, draw a dot 4” beneath it at 30 degrees relative to the dots above. Drill a ½” hole where each dot is located. Make sure that each board looks identical with their holes matching up. Finally, line up two boards 2’ apart and insert the ends of the dowels on the corresponding holes of the boards using wood glue at the tips to make sure they stay in place. Repeat this process until dowels have been placed in all the holes in order to create the frame.

**Figure A2.**
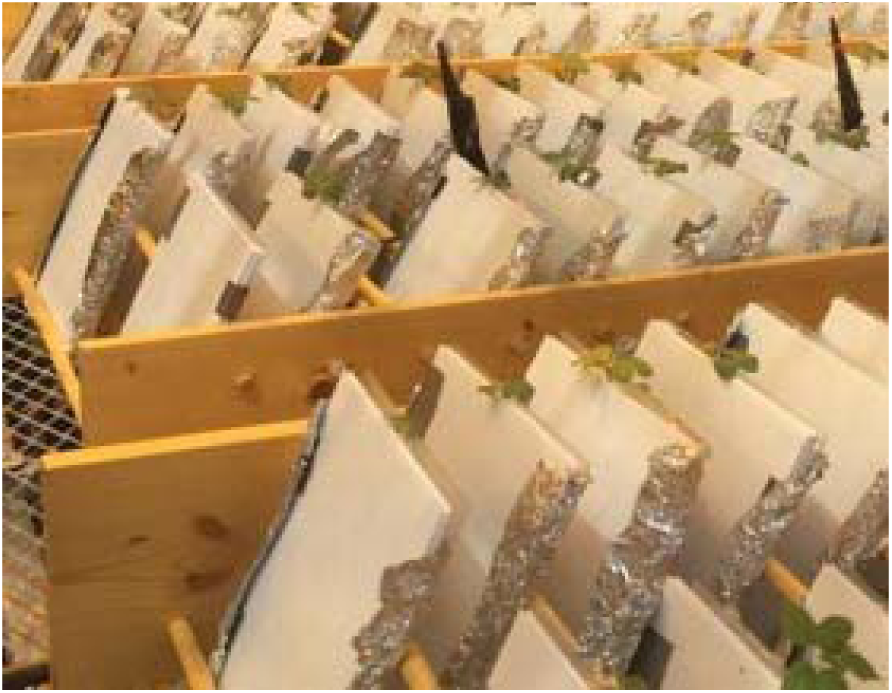
The rhizotron greenhouse experiment. Each rhizotron was maintained at a thirty degree angle with the use of custom built wooden frames (right).

## Appendix B

### Supporting Tables and Figures

**Figure B1.** PCA summarizing root trait space between *I.hederacea* and *I. purpurea* from population PA4 grown in field conditions where 65.2 % and 25.2% of the phenotypic variation is explained by PCA1 and PCA2, respectively. Bar graphs (B & C) show the loading scores of each root trait for PCA1 and PCA2. B) root system size and root system width load heaviest in PCA1, C) and average root angle loads heaviest in PCA2.

**Figure B2.**
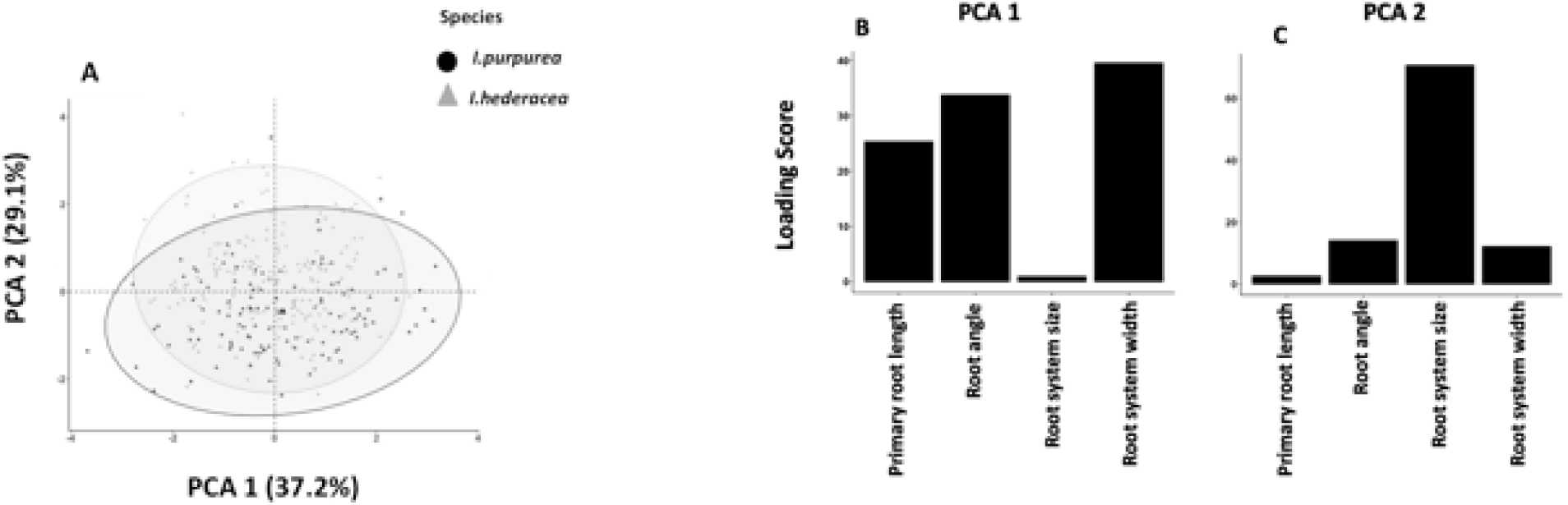
PCA results summarizing the variation of all the root traits measured in the greenhouse experiment (primary root length, root angle, root system size and root system width) between *I. hederacea* and *I. purpurea*. A) The PCA biplot shows the first two PCAs’ and how individuals of *I.purpurea* (black solid circles) and *I.hederacea* (grey solid triangles) vary in this trait space. A) 37.2% and 29.1% of the phenotypic variation is explained by PCA1 and PCA2, respectively, and the overlapping ellipses representing the 95 % confidence intervals of each species indicates high phenotypic similarity between *I.purpurea* and *I.hederacea*. Bar graphs (B & C) show the loading scores that each root trait contributes to PCA1 and PCA2, respectively. B) Root angle, root system width and primary root length load strongest in PCA1, C) whereas root system size loads strongest in PCA 2. PCA 1 can be used as an indicator of root system architecture and PCA 2 can serve as an indicator of root size.

**Table B1.** Influence of competitive treatment on *I. purpurea* and *I. hederacea* root traits when grown in the field. F-statistics showing the effects of block, species and competitive environment, and chi statistics(χ^2^) showing maternal line variation on plant phenotypes. **Degrees of freedom for the linear mixed model the following:** Block: 3; Species: 1; Competition: 2; Maternal line: 1. Significant fixed and random effects (p-value < 0.05) are indicated in bold. Maternal line is abbreviated as ‘ML’.

|  | Block | Species | Competition | ML |
| --- | --- | --- | --- | --- |
| Trait | F | F | F | $\chi^2$ |
| Root system width (cm) | 1.50 | 0.87 | 0.76 | **5.81** |
| Root system size (cm<sup>2</sup>) | **29.91** | 0.04 | 2.33 | 0.02 |
| Root angle (degrees) | 2.54 | 0.06 | 2.04 | 0.07 |
| Seed number | **9.41** | 0.30 | **9.81** | **5.43** |

**Table B2.** Genetic correlation matrix between root traits for *I. purpurea* (above diagonal) and for *I. hederacea* (below diagonal). Pearson correlations were calculated on the family means of non-transformed root traits for each species separately, and across treatments. Significant correlations (p-value < 0.05) are indicated in bold.

|  | Root angle | Root width | Root size |
| --- | --- | --- | --- |
| Root angle | — | -0.01 | 0.1 |
| Root system width | **0.59** | — | **0.85** |
| Root system size | **0.60** | **0.80** | — |

**Table B3.** Influence of interspecific and intraspecific competitive environment on selection gradients (multivariate analyses) in *I. purpurea* and *I. hederacea*. Selection gradients that are significantly different than zero are shown in bold (p-value < 0.05) whereas gradients that are marginally significant (p-value =/< 0.10) shown with an (^). Standard errors of the estimate presented as ±1 SE. Values in bold indicate significant selection gradient.

|  | *I. purpurea* |  | *I. hederacea* |  |  |
| --- | --- | --- | --- | --- | --- |
| Trait | Alone | Competition (Inter) | Alone | Competition (Inter) | Competition (Intra) |
| Root system width (cm) | -0.24 ±0.21 | 0.37 ±0.21 | 0.16 ±0.12 | -0.22 ±0.16 | -0.03 ±0.13 |
| Root system size (cm<sup>2</sup>) | 0.56<sup>^</sup> ±0.29 | -0.15 ±0.22 | -0.09 ±0.13 | 0.07 ±0.11 | 0.03 ±0.17 |
| Root angle (degrees) | 0.02 ±0.10 | -0.09 ±0.08 | 0.01 ±0.08 | **0.23 ±0.09** | -0.27 ±0.24 |

**Table B4.**
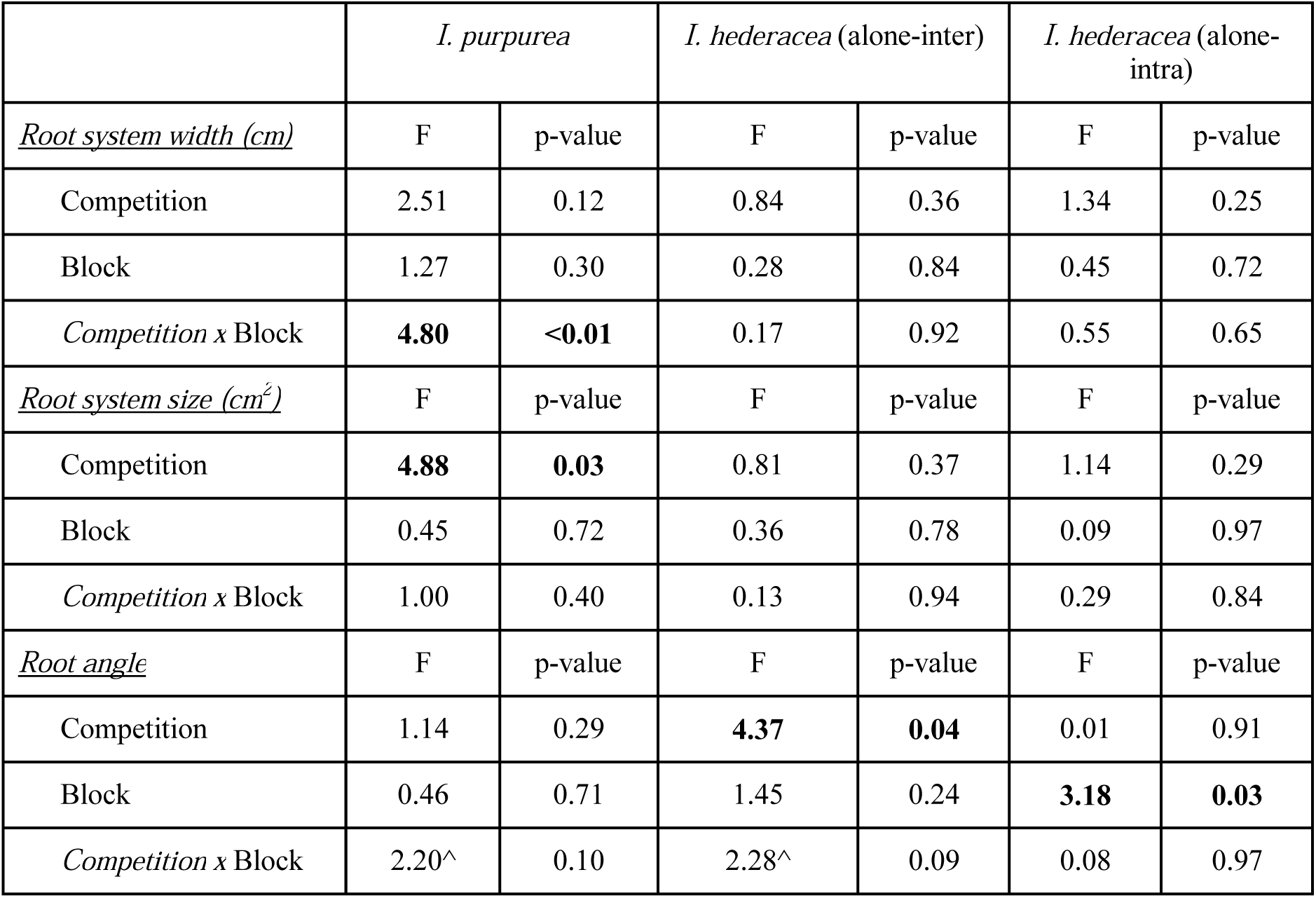
Influence of interspecific competition within *I. purpurea* and interspecific and intraspecific competition in *I. hederacea*, on the pattern of selection acting on specific root raits in these species. F-statistics from the ANCOVA testing the effects of competition, block, and root trait *x* competition on selection gradients are shown. **Degrees of freedom for the linear mixed model are the following:** Competition: 1; Block: 3; Competition *x* Block: 7. Values in bold indicate significant treatment *x* trait, block *x* trait, and treatment *x* block *x* trait interactions whereas effects that are marginally significant (p-value =/< 0.10) shown with an (^).

## Notes

#### Summary of Updates

This version of the manuscripts has been revised to include a new figure, and reduce the word count.

## References

Aarssen, L. W. 1997. High productivity in grassland ecosystems: affected by species diversity or productive species? Oikos 80: 182–183.

Abràmoff, M. D., and P. J. Magalhães. 2004. Image processing with ImageJ. Biophotonics.

Arthur, W. 1982. The evolutionary consequences of interspecific competition. Adv. Ecol. Res. 12:127–187.

Barkley, T. M. 1986. Flora of the Great Plains. University Press of Kansas, Lawrence.

Bartelheimer M, T. Steinlein, W. Beyschlag. 2006. Aggregative root placement: a feature during interspecific competition in inland sand-dune habitats. Plant and Soil 280:101–114.

BassiriRad, H. 2005. Nutrient acquisition by plants: an ecological perspective. Springer science & business media.

Bates, D., M. Mächler, B. Bolker, and S. Walker. 2015. Fitting linear mixed-effects models using lme4. Journal of Statistical Software, Articles 67 (1): 1–48.

Baucom, R. S., S-M Chang, J. M. Kniskern, M. D. Rausher, and J. R. Stinchcombe. 2011. Morning glory as a powerful model in ecological genomics: tracing adaptation through both natural and artificial selection. Heredity 107 (5): 377–85.

Baylis, A.D. 2000. Why glyphosate is a global herbicide: strengths, weaknesses and prospects. Pest Management Science 56 (4): 299–308.

Beebe, S.E., M. Rojas-Pierce, X. Yan, M.W. Blair, F. Pedraza, F. Muñoz, J. Tohme, and J.P. Lynch. 2006. Quantitative trait loci for root architecture traits correlated with phosphorus acquisition in common bean. Crop Science 46: 413–23.

Belter, P.R., and J.F. Cahill Jr. 2015. Disentangling root system responses to neighbours: identification of novel root behavioural strategies. AoB Plants, 7, plv059.

Bright, K. 1998. Geographic variation and natural selection on a leaf shape polymorphism in the ivyleaf morning glory (Ipomoea hederacea). PhD diss. Duke University, Durham, NC.

Briones, O., C. Montaña, and E. Ezcurra. 1996. Competition between three chihuahuan desert species: evidence from plant size-distance relations and root distribution. Journal of Vegetation Science: Official Organ of the International Association for Vegetation Science 7 (3): 453–60.

Brown, W. L., and E. O. Wilson. 1956. Character displacement. Systematic Zoology 5 (2): 49–64.

Burns, Jean H., and Sharon Y. Strauss. 2011. More closely related species are more ecologically similar in an experimental test. Proceedings of the National Academy of Sciences of the United States of America 108 (13): 5302–7.

Cahill, Jr, J.F., and G.G. McNickle, 2011. The behavioral ecology of nutrient foraging by plants. Annual Review of Ecology, Evolution, and Systematics 42 (1): 289–311.

Callaway, R. M., S. C. Pennings, and C. L. Richards. 2003. Phenotypic plasticity and interactions among plants. Ecology.

Callaway, R.M. 2002. The detection of neighbors by plants. Trends in Ecology & Evolution 17 (3): 104–5.

Casper, B.B., and R.B. Jackson. 1997. Plant competition underground. Annual Review of Ecology and Systematics 28 (1): 545–70.

Casper, Brenda B., H.J. Schenk, and R.B. Jackson. 2003. Defining a plant’s belowground zone of influence. Ecology 84 (9): 2313–21.

Colombi, Tino, N. Kirchgessner, C.A. Le Marié, Larry Matthew York, Jonathan P. Lynch, and Andreas Hund. 2015. Next generation shovelomics: set up a tent and REST. Plant and Soil 388 (1): 1–20.

Conner, J.K., R. Franks, and C. Stewart. 2003. Expression of additive genetic variances and covariances for wild radish floral traits: comparison between field and greenhouse environments. Evolution; International Journal of Organic Evolution 57 (3): 487–95.

Day, T., and K.A. Young. 2004. Competitive and facilitative evolutionary diversification. BioScience 54:101–109.

Depuydt, S. 2014. Arguments for and against self and non-self root recognition in plants. Frontiers in Plant Science 5: 614.

Dudley, S. A. 1996. Differing selection on plant physiological traits in response to environmental water availability: a test of adaptive hypotheses. Evolution 50: 92–102.

Dudley, S. A. and A. L. File. 2007. Kin recognition in an annual plant. Biol. Lett. 3: 435–438.

Ehdaie, B., and J.G. Waines. 2008. larger root system increases water–nitrogen uptake and grain yield in bread wheat. In: Appels R et al (eds) 11th international wheat genetics symposium 2008. Sydney University Press, Brisbane, p 659.

Faget, M., N.A. Kerstin, A. Walter, J.M. Herrera, S. Jahnke, U. Schurr, and V.M. Temperton. 2013. Root-root interactions: extending our perspective to be more inclusive of the range of theories in ecology and agriculture using in-vivo analyses. Annals of Botany 112 (2): 253–66.

Falik O, P. Reides, G.M. Novoplansky A. 2003. Self/non-self discrimination in roots. Journal of Ecology 91:525–531.

Fang, Zhou, A.M. Gonzales, M.L. Durbin, Kapua K. T. Meyer, B.H. Miller, K.M. Volz, M.T. Clegg, and P.L. Morrell. 2013. Tracing the geographic origins of weedy Ipomoea purpurea in the southeastern United States. The Journal of Heredity 104 (5): 666–77.

Fang S, R.T. Clark, Y. Zheng, A.S. Iyer-Pascuzzi, J.S. Weitz, L.V. Kochian, H. Edelsbrunner, H. Liao, P.N. Benfey. 2013. Genotypic recognition and spatial responses by rice roots. Proceedings of the National Academy of Sciences of the USA 110:2670 –2675.

Fargione, J., and D. Tilman, 2005. Niche differences in phenology and rooting depth promote coexistence with a dominant c4 bunchgrass. Oecologia 143 (4): 598–606.

Ferguson, L, G., Sancho, M.T. Rutter, and C.T. Murren. 2016. Root architecture, plant size and soil nutrient variation in natural populations of arabidopsis thaliana. Evolutionary Ecology 30 (1): 155–71.

Fitter, A. H., T. R. Stickland, M. L. Harvey, and G. W. Wilson. 1991. Architectural analysis of plant root systems 1. Architectural correlates of exploitation efficiency. The New Phytologist 118 (3): 375–82.

Fitter, A.H. 2002. Characteristics and functions of root systems. In Plant Roots, 49–78. CRC Press.

Fitter, A.H., L. Williamson, B. Linkohr, and O. Leyser. 2002. Root system architecture determines fitness in an arabidopsis mutant in competition for immobile phosphate ions but not for nitrate ions. Proceedings. Biological Sciences / The Royal Society 269 (1504): 2017–22.

Gause, G. F. 1936. The struggle for existence. Soil Science 41 (2): 159.

Gray, A. 1886. Flora of North America. Ivison, Blakeman, Taylor, London.

Hardin, G. 1960. The competitive exclusion principle. Science 131 (3409): 1292–97.

Hickman, J. C. 1993. The Jepson manual: higher plants of California. University of California Press, Berkeley.

Ho, M.D., J.C. Rosas, K.M. Brown, and J.P. Lynch. 2005. Root architectural tradeoffs for water and phosphorus acquisition. Functional Plant Biology: FPB 32 (8): 737–48.

Hodge, A., D. Robinson, B.S. Griffiths and A. H. Fitter. 1999. Why plants bother: root proliferation results in increased nitrogen capture from an organic patch when two grasses compete. Plant, Cell & Environment 22 (7): 811–20.

Huston, M.A. 1997. Hidden treatments in ecological experiments: re-evaluating the ecosystem function of biodiversity. Oecologia 110 (4): 449–60.

Hutchinson, G. Evelyn. 1957. Concluding remarks. Cold Spring Harbor Symposia on Quantitative Biology 22 (January): 415–27.

Kellermeier, F., F. Chardon, and A. Amtmann. 2013. Natural variation of arabidopsis root architecture reveals complementing adaptive strategies to potassium starvation. Plant Physiology 161 (3): 1421–32.

Kuznetsova, Alexandra, P. Brockhoff, and R. Christensen. 2017. lmerTest package: tests in linear mixed effects models. Journal of Statistical Software, Articles 82 (13): 1–26.

Lambers, H., M.W. Shane, M.D. Cramer, S.J. Pearse, and E.J. Veneklaas. 2006. Root structure and functioning for efficient acquisition of phosphorus: matching morphological and physiological traits. Annals of Botany 98 (4): 693–713.

Lande, R., and S.J. Arnold. 1983. The measurement of selection on correlated characters. Evolution 37 (6): 1210–26.

Lê S., J. Josse, F. Husson. 2008 FactoMineR: An R package for multivariate analysis. Journal of Statistical Software, 25(1), 1–18.

Litav M, J.L. Harper. 1967. A method for studying spatial relationships between the root systems of two neighboring plants. Plant and Soil 26:389–392.

Loreau, Michel. 2000. Biodiversity and ecosystem functioning: recent theoretical advances. Oikos 91 (1): 3–17.

Lynch, J. 1995. Root architecture and plant productivity. Plant Physiol. 109: 7–13.

Lynch, J.P., and K.M. Brown. 2001. Topsoil foraging – an architectural adaptation of plants to low phosphorus availability. Plant and Soil 237 (2): 225–37.

Lynch, J.P. 2007. Roots of the second green revolution. Australian Journal of Botany 55 (5): 493–512.

MacArthur, R., and R. Levins. 1967. The limiting similarity, convergence, and divergence of coexisting species. The American Naturalist 101 (921): 377–85.

Manschadi, A.M., J. Christopher, P. deVoil, and G.L. Hammer. 2006. The role of root architectural traits in adaptation of wheat to water-limited environments. Functional Plant Biology: FPB 33 (9): 823–37.

Mauricio, R., and M.D. Rausher. 1997. Experimental manipulation of putative selective agents provides evidence for the role of natural enemies in the evolution of plant defense. Evolution 51 (5): 1435–44.

Mahall, B. E., and R. M. Callaway. 1991. Root Communication among desert shrubs. Proceedings of the National Academy of Sciences of the United States of America 88 (3): 874–76.

Mahler, W. F. 1984. Shinners’ manual of the north central Texas flora. Botanical Research Institute of Texas, Forth Worth.

Menne, M.J., I. Durre, R.S. Vose, B.E. Gleason, and T.G. Houston. 2012. An overview of the global historical climatology network-daily database. Journal of Atmospheric and Oceanic Technology (29): 897–910. doi.10.1175/JTECH-D-11-00103.1.

Mohr, C. 1901. Plant life of Alabama. Brown, Montgomery, AL.

Mueller, K.E., D. Tilman, D.A. Fornara, and S.E. Hobbie. 2013a. Root depth distribution and the diversity—productivity relationship in a long-term grassland experiment. Ecology 94 (4): 787–93.

Paez-Garcia A., C.M. Motes, W.R. Scheible, R. Chen, E.B. Blancaflor, and M.J. Monteros. 2015. Root traits and phenotyping strategies for plant improvement. Plants 4 (2): 334–55.

Pfennig, K. S., and D. W. Pfennig. 2009. Character displacement: ecological and reproductive responses to a common evolutionary problem. The Quarterly Review of Biology.

Poorter, H., and P. Ryser. 2015. The limits to leaf and root plasticity: what is so special about specific root length? The New Phytologist 206 (4): 1188–90.

Rausher, M.D. 1992. The measurement of selection on quantitative traits: biases due to environmental covariances between traits and fitness. Evolution 46 (3): 616–26.

Rubio, G., H. Liao, X. Yan, and J.P. Lynch. 2003. Topsoil foraging and its role in plant competitiveness for phosphorus in common bean. Crop Science 43: 598–607.

Sandhu, N., K. A. Raman, R.O. Torres, A. Audebert, A. Dardou, A. Kumar, and A. Henry. 2016. Rice root architectural plasticity traits and genetic regions for adaptability to variable cultivation and stress conditions. Plant Physiology 171 (4): 2562–76.

Schenk, H. J. 2006. Root competition: beyond resource depletion: root competition: beyond resource depletion. The Journal of Ecology 94 (4): 725–39.

Schluter, D., and J. D. McPhail. 1992. Ecological character displacement and speciation in sticklebacks. The American Naturalist 140 (1): 85–108.

Semchenko M, E.A. John, M.J. Hutchings. 2007. Effects of physical connection and genetic identity of neighbouring ramets on root-placement patterns in two clonal species. New Phytologist 176:644–654.

Silvertown, J. 2014. Plant coexistence and the niche. Trends in Ecology & Evolution 19 (11): 605–11.

Shreve, F., M. A. Chrysler, F. H. Blodgett, and F. W. Besley. 1910. The plant life of Maryland. Johns Hopkins University Press, Baltimore.

Smith, R.A., and M.D. Rausher. 2008. Experimental evidence that selection favors character displacement in the ivyleaf morning glory. The American Naturalist 171 (1): 1–9.

Strausbaugh, P. D., and E. L. Core. 1964. Flora of West Virginia. West Virginia University Bulletin, Morgantown.

Stevens, W. C. 1948. Kansas wildflowers. University of Kansas Press, Lawrence.

Svačina, P., T. Středa, and O. Chloupek. 2014. Uncommon selection by root system size increases barley yield. Agronomy for Sustainable Development 34 (2): 545–51.

Tilman, D., C. L. Lehman, and K. T. Thomson. 1997. Plant diversity and ecosystem productivity: theoretical considerations. Proceedings of the National Academy of Sciences of the United States of America 94 (5): 1857–61.

Tilman, D., P. B. Reich, J. Knops, D. Wedin, T. Mielke, and C. Lehman. 2001. Diversity and productivity in a long-term grassland experiment. Science 294 (5543): 843–45.

Uga, Y., K. Sugimoto, S. Ogawa, J. Rane, M. Ishitani, N. Hara, Y. Kitomi, et al. 2013. “Control of root system architecture by DEEPER ROOTING 1 increases rice yield under drought conditions.” Nature Genetics 45 (9): 1097–1102.

Wade, M.J. and S. Kalisz. 1990. The causes of natural selection. Evolution 44 (8): 1947–55.

Wasson, A. P., R. A. Richards, R. Chatrath, S. C. Misra, S. V. Sai Prasad, G. J. Rebetzke, J. A. Kirkegaard, J. Christopher, and M. Watt. 2012. Traits and selection strategies to improve root systems and water uptake in water-limited wheat crops. Journal of Experimental Botany 63 (9): 3485–98.

Wunderlin, R. P. 1982. Guide to the vascular plants of central Florida. University Presses of Florida, Tampa.

York, L.M., T. Galindo-Castañeda, J.R. Schussler, and J.P. Lynch. 2015. Evolution of us maize (zea mays l.) Root architectural and anatomical phenes over the past 100 years corresponds to increased tolerance of nitrogen stress. Journal of Experimental Botany 66 (8): 2347–58.

